# Genomic insights into *Plasmodium vivax* population structure and diversity in central Africa

**DOI:** 10.1101/2022.12.16.520826

**Authors:** Valerie Gartner, Benjamin D. Redelings, Claudia Gaither, Jonathan B. Parr, Albert Kalonji, Fernandine Phanzu, Nicholas F. Brazeau, Jonathan J. Juliano, Gregory A. Wray

## Abstract

*Plasmodium vivax* malaria has not traditionally been a major concern in central Africa given the high prevalence of the human Duffy-negative phenotype that is believed to prevent infection. Increasing reports of asymptomatic and symptomatic infections in Duffy-negative individuals throughout Africa raise the possibility that *P. vivax* is evolving to evade host resistance, but there are few parasite samples with genomic data available from this part of the world. In this study, we perform whole genome sequencing of a new *P. vivax* isolate from the Democratic Republic of the Congo (DRC) and assess how this central African isolate fits into the global context of this species. We use population genomics methods to show that *P. vivax* from DRC is similar to other African parasite populations and is not closely related to the non-human primate parasite *P. vivax*-like.

**Significance:** The second most common malaria species to infect humans, *Plasmodium vivax*, is not considered a major threat to human health in central Africa because people in this region frequently have a genetic variant that prevents the *P. vivax* species from being able to cause illness. Recent research shows that *P. vivax* can be found in individuals who should be immune, but there is insufficient data to understand why. Our study investigates the genome of one *P. vivax* sample collected from central Africa to show that the DRC population is closely related to other *P*.*vivax* populations in Africa.

## Introduction

The widespread fixation of the Duffy-negative phenotype in the human population in Sub-Saharan Africa, which provides immunity from *Plasmodium vivax*, is one of the most remarkable cases of natural selection documented in human populations (***Hamblin and Di Rienzo, 2000; Hamblin et al., 2002; Kwiatkowski, 2005***). The Duffy-negative phenotype occurs in humans with two copies of a silencing mutation in the promoter region of the Duffy Antigen Receptor for Chemokines (DARC) gene, resulting in the absence of receptor expression exclusively in erythrocytes necessary for the progression of the *P. vivax* life cycle (***Parasol et al., 1998; Miller et al., 1976***). Despite this, there are an increasing number of reports of asymptomatic and symptomatic *P. vivax* infections in people with the Duffy-negative mutation suggesting that *P. vivax* persists in central Africa at low levels in people with the Duffy-negative resistance allele (***Brazeau et al., 2021; Russo et al., 2017; Motshoge et al., 2016; Ryan et al., 2006; Mendes et al., 2011***).

An alternate explanation for the persistence of *P. vivax* in central Africa comes from the recent discovery of a closely related parasite species that infects non-human primates, *Plasmodium vivax*-like in Western Africa (***Liu et al., 2014; Loy et al., 2018; Gilabert et al., 2018***). Though there is only one confirmed report of *P. vivax*-like infecting humans, a study using *P. vivax*-like recombinant binding proteins did not reveal species-specific barriers to erythrocyte invasion of human, gorilla, or chimpanzee red blood cells, suggesting *P. vivax*-like likely is able to infect humans (***Prugnolle et al., 2013; Loy et al., 2018***).

A third possible explanation for the presence of *P. vivax* in central Africa despite human resistance alleles might be that *P. vivax* is adapting to overcome the Duffy negative resistance allele, which would be a serious concern for malaria elimination efforts in central Africa. Genomics can potentially aid in understanding the source of these infections in central Africa, however none of the seventy-seven publicly available African *P. vivax* genomes are from regions with high levels of Duffy negativity except for three samples from Uganda. Importantly, these Ugandan samples were collected from people of unknown Duffy status after returning to the United Kingdom (***Gunalan et al., 2018; Benavente et al., 2021***).

In this study, we perform whole genome sequencing of a new *P. vivax* isolate from the Eastern region of Democratic Republic of the Congo (DRC) and assess how *P. vivax* from central Africa fits into the global context of this pathogen. Though the original study design excludes the possibility of genotyping the human host of this *P. vivax* sample, the patient had no known travel history and resides in a region where the Duffy-negative phenotype frequency is at or above 80% (***Howes et al., 2011***), thus this patient has a high chance of having the Duffy-negative phenotype. We confirm the presence of a *P. vivax* population in central Africa that is not closely related to the ape-infection *P. vivax*-like species. Further, we investigate whether this sample has duplications of the Duffy binding ligand genes PvDBP and PvDPB2 (also referred to as EBP and EBP2) which might potentially enable *P. vivax* to evade host immunity. Though Copy Number Variation of these genes is not conclusively linked to *P. vivax* infection of Duffy-negative individuals (***Lo et al., 2019, 2021***), both genes contain the Duffy Binding Protein II domain, one of the foremost vaccine target candidates (***Roesch et al., 2018***). We show a duplication of the gene PvDBP in the DRC *P. vivax* sample and a single copy of PvDBP2.

## Results

### *P. vivax* from DRC falls within African diversity in context of global population structure

To determine where this new central African sample fits within global *P. vivax* populations, we first performed principal components analysis (PCA) from 696 *P. vivax* samples using only biallelic SNPs (excluding hyper-variable regions defined by (***Pearson et al., 2016; Auburn et al., 2016***)). The global PCA analysis in Figure 1A shows a population structure as defined by geography is reproduced by the first two principal components, as has been reported previously (***Daron et al., 2021; Benavente et al., 2021***). Three main sub-populations are formed: 1. samples from the Americas, 2. African and South Asian samples, and 3. East Asian and Southeast Asian samples. Within the cluster of African and South Asian samples, the new DRC sample is most similar to those from Uganda and Madagascar in their position alongside South Asian *P. vivax*.

**Figure 1.**
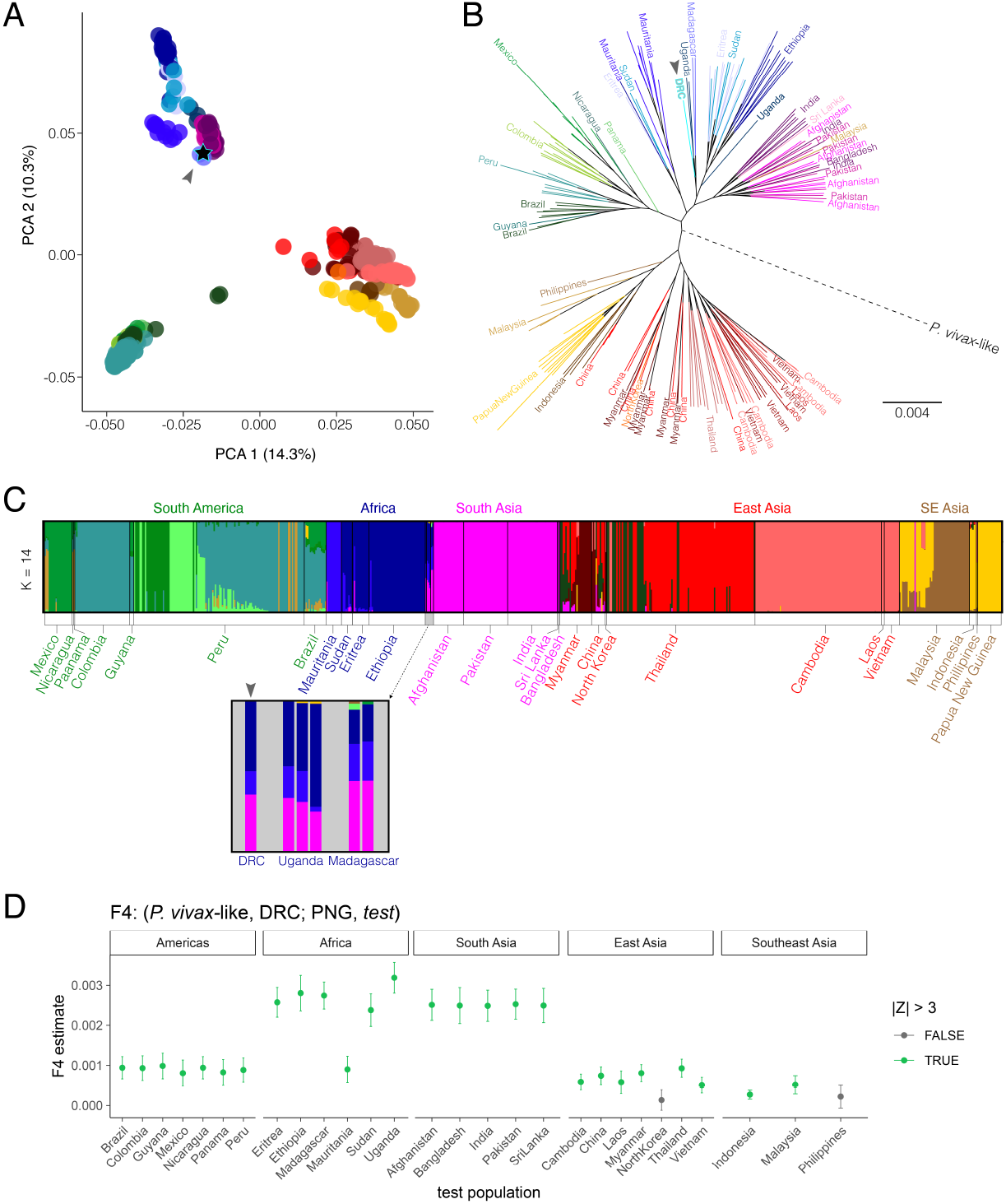
DRC P. vivax falls within African variation in global species context. DRC sample indicated by arrowhead in each panel. **A** Principal components analysis of global genetic diversity reveals *P. vivax* from the DRC grouped with African and South Asian populations. Thefirst principal component explains 14.3% of variation in the data and the second principal component represents 10.3% of variation. **B** Maximum Likelihood tree shows *P. vivax* from the DRC clusters with Uganda and Madagascar. Tree constructed using whole-genome SNP data and rooted using two *P. vivax*-like sequences (red dotted line). Continent colors: Americas (green), Africa (blue), South Asia (pink), East Asia (red), Southeast Asia (brown). Tree constructed using IQtree with the GTR+ASC model to account for ascertainment bias in SNP data. **C** Admixture analysis of global population structure indicates *P. vivax* ancestry is geographically structured. Each vertical bar represents the proportion of genetic ancestry belonging to one individual *P. vivax* sample for each simulated population size (K). Population size K=14 is most-supported by Cross Validation Error. **D** F4 statistics calculated using Admixtools2 using the relationship (*P. vivax*-like, DRC; Papua New Guinea (PNG), *test*). Higher F4 estimates indicate DRC has a closer relationship with the test population than it does to the PNG population. Error bars indicate ±3 SE.

To further understand how this new DRC *P. vivax* sample relates to other global populations, we constructed a maximum likelihood tree from 349,353 SNPs across the genome for no more than 10 samples per country. Figure 1B shows the DRC sample clusters within African *P. vivax* variation, and again clusters most closely with Uganda and Madagascar samples. *P. vivax*-like samples were used to root this tree, and notably, as in ***Daron et al. (2021)***, the root for our tree is located centrally in the *P. vivax* tree and not inside African variation, which might be expected if this sample represented an ancestral source population of *P. vivax* in humans.

Though clustering of genetic variation is expected due to geographic separation, PCA and phylo-genetic trees do not quantify how much genetic ancestry is shared across geographic populations. To determine the fraction of shared ancestry across subgroups, we assessed the same SNP data set used in Figure 1A-B through Admixture analysis. We modeled *P. vivax* ancestry proportions for population sizes of 2 through 17 and found that a population size of K=14 was supported based on mean Cross Validation Error value. Figure 1C shows the global ancestry proportions of *P. vivax* when modeled at K=14. Ancestry proportions for all calculated K values is shown in Supplementary Figure 3. Additionally, F4 statistics (***Patterson et al., 2012; Maier et al., 2022)*** were calculated assess the correlation in allele frequencies of the DRC sample with other *P. vivax* populations around the world. We use the form (*P. vivax*-like, DRC; Papua New Guinea, Y), where *P. vivax*-like is the outgroup species, shown in figure 1D. Higher F4 estimates indicate the DRC sample has more gene flow with the test population than it does with samples from Papua New Guinea (PNG). All test populations except for North Korea and the Philippines resulted in a significant absolute Z score (|Z| > 3). F4 estimates and related data are available in Supplemental table 4.

### *P. vivax* from DRC has similar levels of population diversity as other African populations

In order to explore *P. vivax* population diversity despite having only a single sample from the DRC, we looked at the number of private alleles in each country. Private alleles are variants present in one population and in none of the others, making them unique to a population. Table S1 shows the full set of summary statistics calculated for all countries. When normalizing the private allele count by dividing by the number of samples, as shown in Figure S1B, the DRC sample had a similar amount of variation as other African populations despite having a low absolute private allele count (Figure S1A). The genome-wide within-population diversity value, π, calculated for *P. vivax* from different sub-regions shown in Figure S2 indicates that when combined, all African samples have similar genome-wide diversity as populations in Asia. *P. vivax* nucleotide diversity of central African samples (DRC and Uganda) is similar to but slightly lower than that of East African (Ethiopia, Eritrea, and Sudan).

### *P. vivax* in central Africa is distinct from *P. vivax*-like

To assess the relatedness of *P. vivax* in humans in central Africa to the *P. vivax*-like malaria species found in non-human primates, we constructed a maximum likelihood tree including publicly available *P. vivax*-like genome sequences mapped to the PvP01 reference genome. Figure 2 shows that the *P. vivax* sample from the DRC clusters with other African *P. vivax* samples, while all *P. vivax*-like samples are separate from *P. vivax* populations. Additionally, the longer branch lengths for the *P. vivax*-like samples in figure 2 illustrate the higher level of diversity in this species than is found in any population of the human-infecting *P. vivax*.

**Figure 2.**
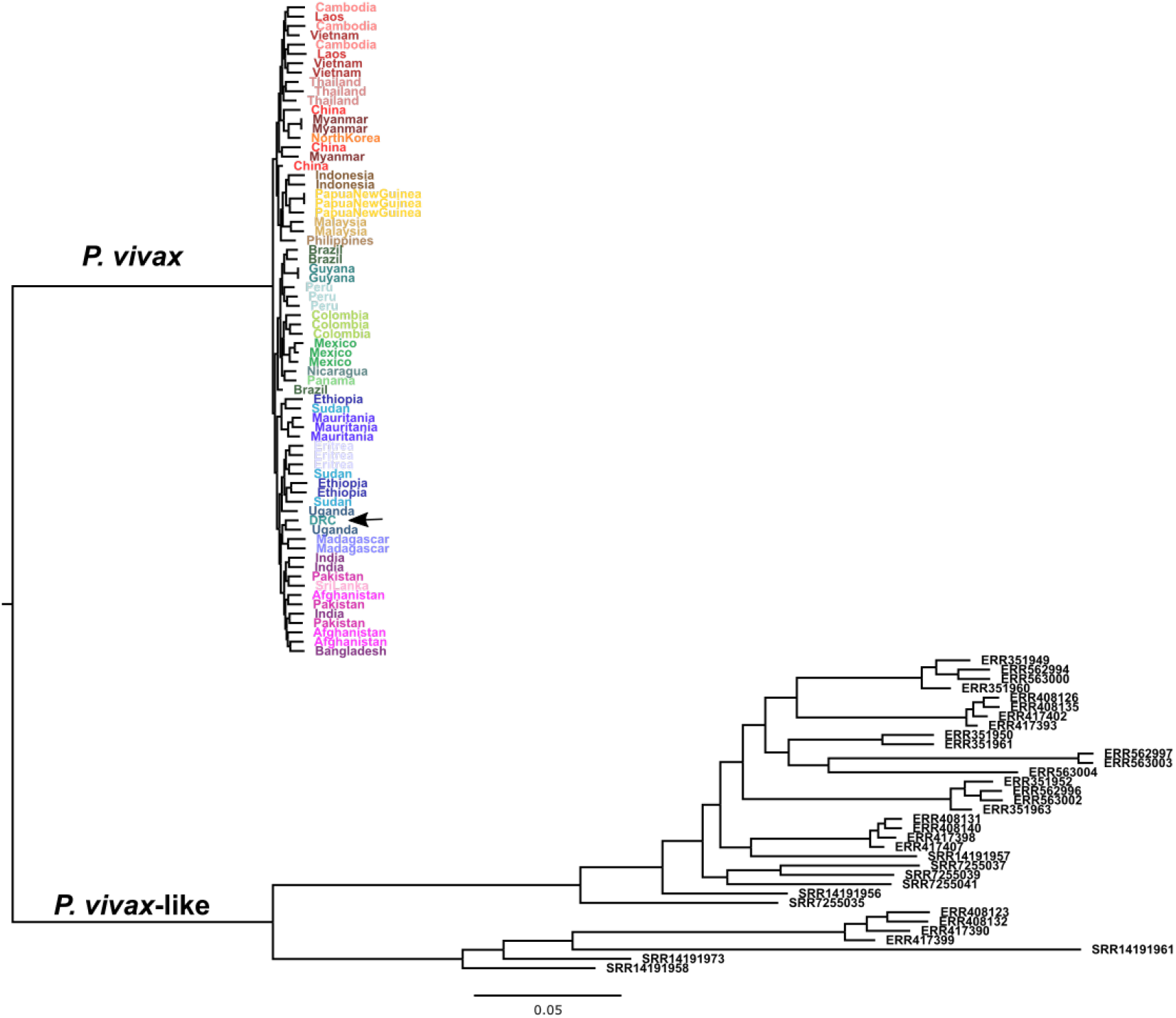
Maximum Likelihood tree shows DRC *P. vivax* sample branches with *P. vivax* populations and not *P. vivax*-like. Maximum Likelihood tree of nuclear genome SNP data shows that *P. vivax* from the DRC does not branch with *P. vivax*-like samples (black tip labels). The DRC sample, indicated here with an arrow, clusters with other African samples.

### Copy number variation is present in binding proteins

Copy Number Variation (CNV) in certain binding proteins is potentially important for pathogenesis of *P. vivax* in Duffy-negative individuals (***Gunalan et al., 2018)***. We used BAM files aligned to PvP01 with optical duplicates removed to compare read depth within the gene region to coverage in the region 10 Kb upstream and downstream of the coding region for several genes related to erythro-cyte binding and invasion: PvDBP, PvDBP2, PvRBP1a, PvRBP1b, PvRBP2a, PvRBP2b, and PvRBP2c based on results from ***Gunalan et al. (2018)***. In the DRC *P. vivax* sample, only one gene, PvDBP, had evidence of a potential gene duplication (Supplemental table 2). Lumpy was used to determine the number of paired-end and split reads that support a duplication in PvDBP, which showed evidence of a duplication of 8,216 base pairs in length at chr6: 980,472-988,688 with 292 paired-end reads and 419 split reads supporting the structural variant. The ratio of the coverage for the duplicated PvDBP region compared to the surrounding intergenic region was 2.47. Based on the IGV pileup view in Figure 3, there appears to be two distinct copies of this gene being mapped to the single PvDBP reference annotation. We note that the region of higher read depth extends into the intergenic regions on either side of the gene annotation for PvDBP, consistent with the duplication type first reported in Malagasy samples (***Menard et al., 2013; Hostetler et al., 2016)***.

**Figure 3.**
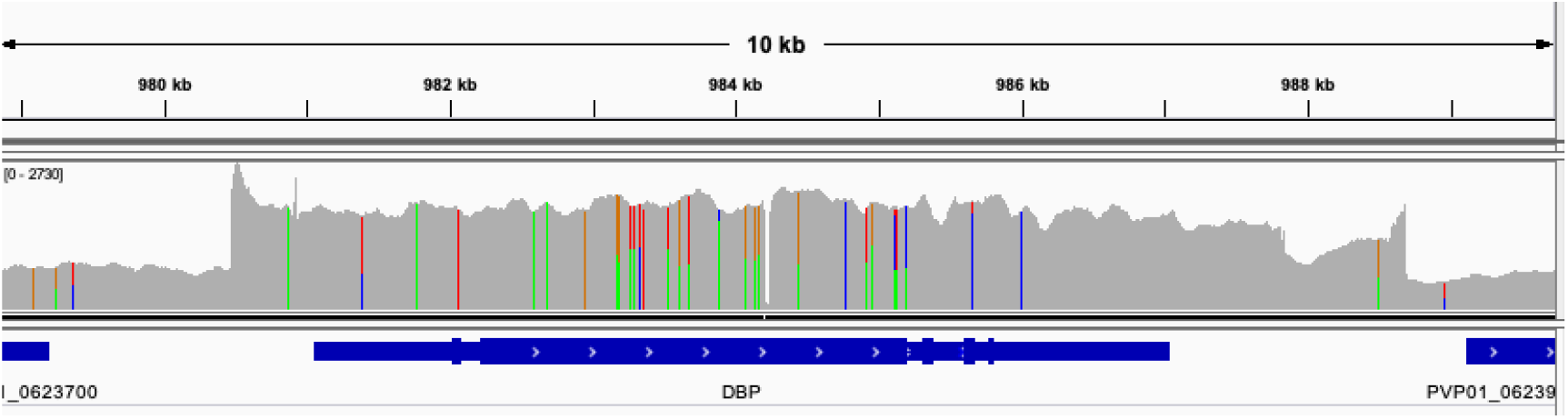
Read Pileup image of duplication of PvDBP in *P. vivax* from DRC. IGV view of the read depth for PvDBP in the DRC *P. vivax* sample where genome coordinate is on the X-axis and read depth is on the Y-axis. This shows an increase in read depth around PvDBP indicating a duplication, and the different variants present along this region suggest two distinct haplotypes.

## Discussion

Despite the publication of recent studies on *P. vivax* diversity that include African samples (***Chan et al., 2012; Menard et al., 2013; Auburn et al., 2019; Twohig et al., 2019; Daron et al., 2021)***, there is still very little known about this pathogen in central Africa. Our analyses of one new *P. vivax* genome collected from the Idjwi island of Lake Kivu in DRC show that this sample falls within the scope of African parasite diversity and is distinct from *P. vivax*-like samples. This suggests that an endemic *P. vivax* population is present in central Africa, as previously proposed by ***Brazeau et al. (2021)***.

Our results shown in Figure 1 suggest the DRC *P. vivax* sample is most like those from Uganda and Madagascar. While the similarity between *P. vivax* from eastern DRC and Uganda is not surprising, it is interesting that *P. vivax* populations in the DRC, Uganda, and Madagascar all share ancestry with South Asian samples, as shown in Figure 1C and D, but no measurable ancestry from Southeast Asian *P. vivax* populations despite a well-documented history of Austronesian human migration into this region (***Anderson et al., 2018; Brucato et al., 2018, 2019)***.

The phylogenetic tree in Figure 1B shows that this new *P. vivax* sample clusters with other African samples and not with any *P. vivax*-like samples that have been sequenced previously, suggesting that there is a *P. vivax* population in the DRC separate from potential zoonosis from an animal reservoir. However, all publicly available *P. vivax*-like sequences to date have been collected from animals in countries on the West coast of Africa. Further sampling of both humans and non-human primates throughout the broad geography of central Africa is needed to determine whether there truly is no transfer of parasites across species, as *in vitro* studies indicate that there is little host specificity of *P. vivax*-like (***Loy et al., 2018)***.

Our analyses replicate previous studies showing that global *P. vivax* populations are distinct from each other based on geographic distance, and most sharing of haplotype backgrounds occurs within geographic regions and only rarely across geographic borders. This geographic separation of ancestral groups, along with the summary statistics calculated in Supplementary table 1 and illustrated in Supplementary Figures S1 and S2, possibly indicate that DRC has a comparable *P. vivax* population size relative to Ethiopia and Uganda, though this interpretation is greatly limited by the reductive nature of genome-wide summary statistics.

Though the evidence linking copy number of Duffy binding ligand genes with *P. vivax* infection of Duffy-negative individuals is not conclusive (***Lo et al., 2019, 2021)***, it remains a subject of concern, especially since the Duffy Binding Protein-II domain is one of the foremost vaccine target candidates (***Roesch et al., 2018)***. We show that this *P. vivax* sample from the DRC has a duplication in PvDBP relative to the PvP01 reference genome that corresponds with the longer Malagasy-type duplication, as opposed the shorter PvDBP duplication first detected in Cambodian samples (***Hostetler et al., 2016)***. Duplications in PvDBP may play a role in Duffy-independent mechanisms of infection in Duffy-negative individuals and should be considered in future studies (***Lo et al., 2021)***.

Our findings are largely limited by the uncertainty of whether this single sample is representative of the larger population of *P. vivax* in central Africa. This *P. vivax* sample was collected from an individual with no known travel history in a region with an estimated 98% homozygosity for Duffy-negativity (***Howes et al., 2011)***, however as the patient’s Duffy genotype was not collected in the study, caution should be exercised until future studies can provide further context.

It has become clear from epidemiological studies that *P. vivax* is much more common in central Africa than previously thought (***Ryan et al., 2006; Twohig et al., 2019; Mendes et al., 2011; Brazeau et al., 2021)***. Generating genomes from these infections however is difficult, as most of them are extremely low parasite densities (***Brazeau et al., 2021)***. Thus, they are not amenable to whole genome sequencing. To increase our understanding of this parasite in Africa, the research community needs to continue to try to identify samples amenable to analysis and deposit them for community use. Intentional sampling across Africa would further contextualize this sample within African *P. vivax* diversity and shed light on the mechanisms of infection in Duffy negative individuals.

## Methods

### Genome Sequencing of *P. vivax* sample from the Democratic Republic of Congo

One *P. vivax* sample was collected from Idjwi, DRC (***Parr et al., 2021)***. The patient was an 11-year-old with reported fever, diarrhea, and headache who tested positive for *P. vivax* via 18s qPCR assay as previously described (***Brazeau et al., 2021)*** at 957 parasites per μL. The patient tested negative for *P. falciparum* via rapid HRP2 test and real time PCR. Due to the original study design, the patient’s Duffy genotype was not assessed. Travel history was not taken.

DNA from three 6mm punches from a dried blood spot were extracted using Chelex-Tween as previously described (***Topazian et al., 2020)***. *P. vivax* infection was confirmed using a Taqman real time PCR assay (***Brazeau et al., 2021)***. *Plasmodium* DNA was enriched from human DNA using a custom Twist hybrid capture array and in-house pipeline (Twist Biosciences, San Francisco, CA, USA). The array was designed by single tiling of the PvP01 genome with baits complementary to human removed. Capture and library preparation were completed per manufacturer’s instructions. Sequencing was completed on a NovaSeq 6000 at the University of North Carolina High Throughput Sequencing Facility.

### Genomic Data Processing

1,408 FASTQ files for *P. vivax* with metadata about the geographic location from which they were sampled were downloaded from the Sequence Read Archive (***Kodama et al., 2012)***. BAM files were created using bwa mem (***Li and Durbin, 2009)*** to align short reads to the PvP01 reference genome (***Auburn et al., 2016)***. Picard MarkDuplicates version 2.18.15 (***Broad Institute, 2019)*** was used to remove optical duplicates, and variants underwent hard filtering using the Genome Analysis Toolkit (GATK) HaplotypeCaller version 3.8.1, followed by joint calling (***Auwera and O’Connor, 2020)***. To avoid confounding analyses with *P. vivax* samples made up of more than one haplotype background (i.e. a multiplicity of infection greater than one), samples were filtered out based on haplotype number estimates generated by Octopus (***Cooke et al., 2018)***. Sample accessions and location metadata of this final sample set of 696 *P. vivax* samples, one being the new DRC genome sequenced in this paper, are available via the project GitHub repo: https://github.com/vlrieg/DRC_vivax/blob/main/sample_info/metadata_table.csv.

### Population Genetics Analysis

Analyses were performed only on assembled chromosomes as defined by the PvP01 reference genome (***Auburn et al., 2016)***. Hyper-variable regions determined in ***Pearson et al. (2016)*** were converted to PvP01 coordinates using the alignment-smc tool in bali-phy with the translate-mask option (***Suchard and Redelings, 2006)***, then removed from the data set using bedtools version 2.25.0 (***Quinlan and Hall, 2010)*** and VCFtools version 0.1.15 (***Danecek et al., 2011)***. The data were reduced to biallelic Single Nucleotide Polymorphisms (SNPs) only, and LD pruning was performed using PLINK to obtain unlinked singletons and variants from the data set as previously described (***Daron et al., 2021; Benavente et al., 2021)***, resulting in 94,083 SNPs. Principal Component Analysis was performed using Plink version 1.9 (***Purcell et al., 2007)*** and plotted in R using ggplot2 (***Wickham, 2016)***. Admixture analysis was performed using Admixture software version 1.3.0 (***Alexander et al., 2009)*** via admixturePipeline version 2.0.2 (***Mussmann et al., 2020)***. The resulting Q matrices were visualized using Pong version 1.5 (***Behr et al., 2016)***. Cross validation error supports K=14 populations. F4 statistics were calculated on 467,205 biallelic SNPs that had no more than 5% missing sites from 696 *P. vivax* and 56 *P. vivax*-like samples using Admixtools2 version 2.0.0 (***Maier et al., 2022)***. Summary statistics were computed on 696 *P. vivax* samples. π was computed on biallelic SNPs using Pixy version 1.2.4.beta1 (***Korunes and Samuk, 2021)***.

### Phylogenetics Analysis

Phylogenies of global and African subsamples of *P. vivax* were made using IQtree version 1.6.12 for Linux 64-bit (***Minh et al., 2020; Hoang et al., 2018)***. Trees were constructed using biallelic SNPs from PvP01-defined nuclear chromosomes except for hypervariable regions. Phylogenies were inferred using the GTR+ASC model to account for ascertainment bias. Trees were rooted using two *P. vivax*-like samples from two recent studies of this closely related species (***Gilabert et al., 2018; Loy et al., 2018)***. Trees were visualized using FigTree version 1.4.4 (http://tree.bio.ed.ac.uk/software/figtree/) and modified with Adobe Illustrator.

### Duffy Binding Gene Copy Number Variation Analysis

The read depths of important genes related to *P. vivax* pathogenesis were investigated by extracting the genomic regions from the BAM file (after removing optical duplicates) using Samtools version 1.3.1 (***Danecek et al., 2021)*** and visualized in IGV version 2.4.14 (***Robinson et al., 2011)***. Genomic coverage as defined by read depth was calculated for PvDBP using bedtools version 2.25.0 (***Quinlan and Hall, 2010)***. Breakpoint evidence to support a duplication of PvDBP was estimated using Lumpy version 0.2.13 (***Layer et al., 2014)***.

## Acknowledgment

This work was supported by: the National Institutes of Health [R01TW010870 and K24AI134990 to J.J.J.], the North Carolina Biotechnology Center support for high-performance computing facility [2016-IDG-1013, 2020-IIG-2109], and the Global Fund to Fight AIDS, Tuberculosis, and Malaria. Thanks to Krista Pipho for feedback on early drafts of this manuscript.

## Author contributions

**Conceptualization:** V.G., B.D.R., N.F.B., J.J.J., G.W. **Sample Collection and Sequencing:** J.B.P., C.G., A.K., F.P., J.J.J. **Conducted Analyses:** V.G. **Advised on Analyses:** B.D.R., N.F.B., J.J.J., G.W. **Aided in Manuscript Preparation:** V.G., B.D.R., C.G., J.B.P, A.K., F.P., N.F.B., J.J.J., G.W.

## Data Availability

*P. vivax* WGS data from DRC available under BioProject accession: PRJNA909777. Accession numbers for previously published data used in this study are available on the project GitHub repository.

