## Supplemental Material for "Genomic insights into *Plasmodium vivax* population structure and diversity in central Africa"

### Supplementary Material

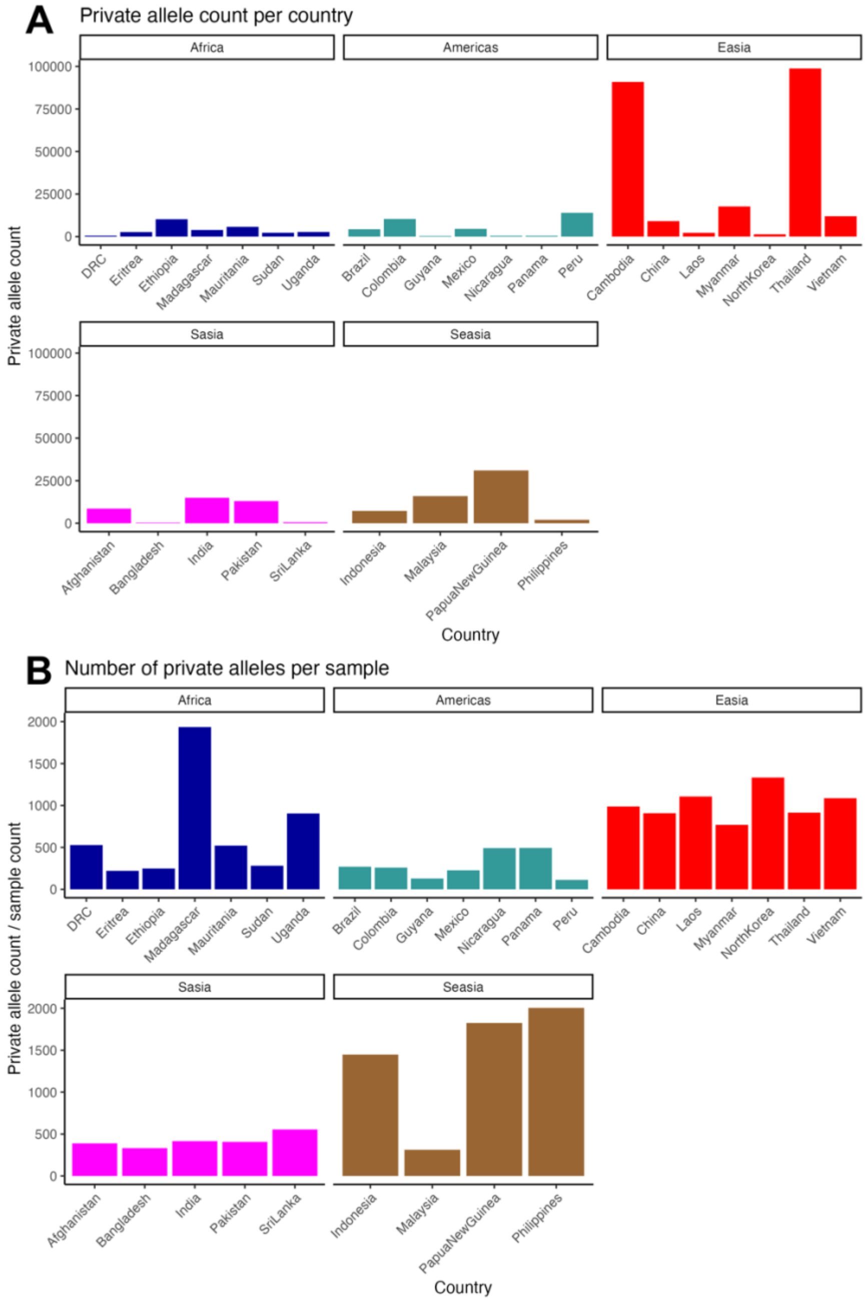


**Supplemental Figure 1. *P. vivax* genome private alleles as a measure of population variation, separated by continent.**

**A.** Absolute count of private alleles for *P. vivax* in each country. *P. vivax* from the DRC has relatively few SNPs that are unique to this population (528 SNPs).

**B.** Private allele count is normalized by dividing by the number of samples in the population. This figure indicates that P. vivax in DRC has a similar private allele count to other African populations, except for Madagascar, once the count is adjusted for the number of samples.

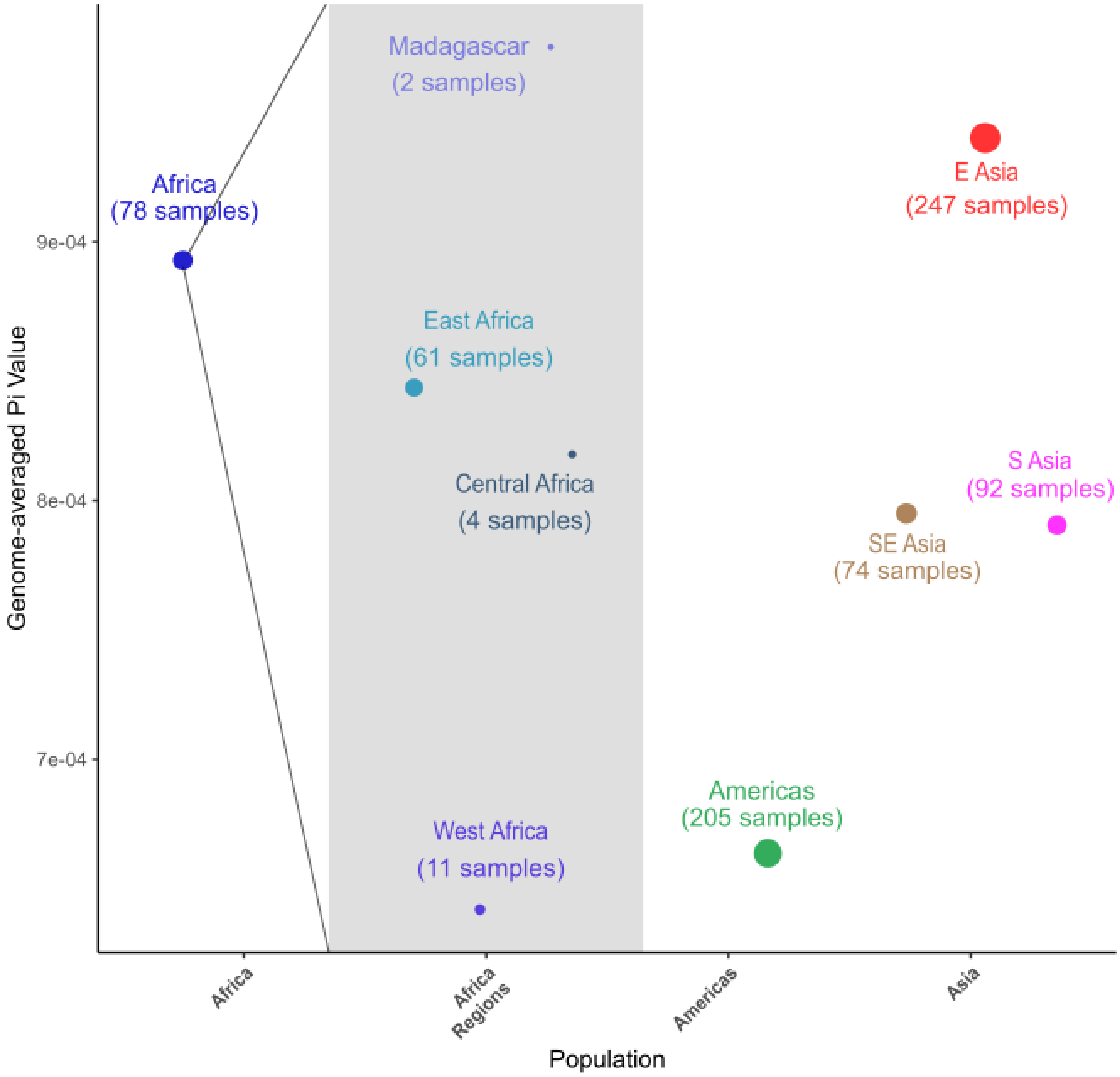


**Supplemental Figure 2. Genome-wide Nucleotide Diversity within Africa**
Genome-wide average nucleotide diversity displayed by continent and by sub-region. African *P. vivax* samples grouped together have similar genome-wide diversity as populations in Asia. Within Africa, nucleotide diversity of central African samples (DRC and Uganda) is similar but slightly lower than that of East African (Ethiopia, Eritrea, and Sudan). (Pi values: Africa = 8.9267e-4; Americas = 6.6364e-4; East Asia = 9.4076e-4; South Asia = 7.9435 e-4; Southeast Asia = 7.9125e-4)

| Region | Country | Number of Private Alleles | Sample Count | Segregating Sites | Private Alleles Per Sample |
|---|---|---|---|---|---|
| Africa | Democratic Republic of Congo | 528 | 1 | 20755 | 528 |
| Africa | Eritrea | 2648 | 12 | 49750 | 220.666667 |
| Africa | Ethiopia | 10195 | 41 | 82966 | 248.658537 |
| Africa | Madagascar | 3871 | 2 | 18169 | 1935.5 |
| Africa | Mauritania | 5739 | 11 | 38829 | 521.727273 |
| Africa | Sudan | 2251 | 8 | 43634 | 281.375 |
| Africa | Uganda | 2715 | 3 | 23558 | 905 |
| Americas | Brazil | 4312 | 16 | 39158 | 269.5 |
| Americas | Colombia | 10362 | 40 | 61993 | 259.05 |
| Americas | Guyana | 388 | 3 | 294 | 129.333333 |
| Americas | Mexico | 4536 | 20 | 33466 | 226.8 |
| Americas | Nicaragua | 492 | 1 | 19040 | 492 |
| Americas | Panama | 494 | 1 | 18376 | 494 |
| Americas | Peru | 13974 | 124 | 66408 | 112.693548 |
| EAsia | Cambodia | 90890 | 92 | 215306 | 987.934783 |
| EAsia | China | 9077 | 10 | 57907 | 907.7 |
| EAsia | Laos | 2214 | 2 | 19180 | 1107 |
| EAsia | Myanmar | 17695 | 23 | 82611 | 769.347826 |
| EAsia | North Korea | 1333 | 1 | 19774 | 1333 |
| EAsia | Thailand | 98794 | 108 | 238360 | 914.759259 |
| EAsia | Vietnam | 11956 | 11 | 67515 | 1086.90909 |
| SAsia | Afghanistan | 8569 | 22 | 66479 | 389.5 |
| SAsia | Bangladesh | 332 | 1 | 20728 | 332 |
| SAsia | India | 14966 | 36 | 92994 | 415.722222 |
| SAsia | Pakistan | 13030 | 32 | 79154 | 407.1875 |
| SAsia | Sri Lanka | 555 | 1 | 22072 | 555 |
| SEAsia | Indonesia | 7239 | 5 | 38468 | 1447.8 |
| SEAsia | Malaysia | 15943 | 51 | 66571 | 312.607843 |
| SEAsia | Papua New Guinea | 31024 | 17 | 74267 | 1824.94118 |
| SEAsia | Philippines | 2005 | 1 | 14277 | 2005 |

**Supplemental Table 1.** *P. vivax* population diversity summary statistics, calculated across 1Kb- long windows along the genome, excluding hyper-variable sites. Private alleles are the number of SNPs unique to that population; segregating sites are the sites that differ from PvP01 reference genome and which are not present at 100% frequency within the population.

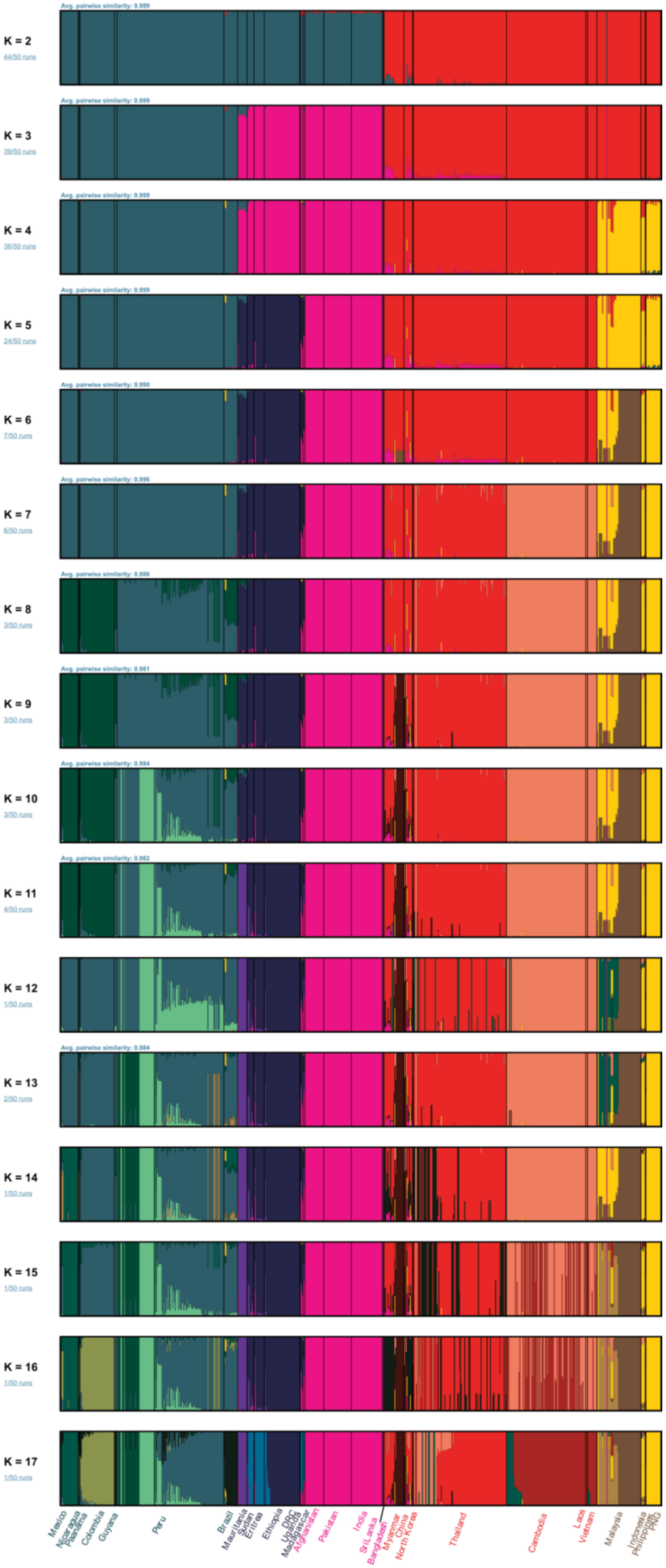


**Supplemental Figure 3. Admixture analysis results for all population sizes** Results of admixture analysis performed for putative population sizes of 2 through 17 (K=2 through K=17). Populations are primarily separated by geography, with little shared ancestry between regions.

| Gene | Chr | Upstream Coordinates | Gene Coordinates | Downstream Coordinates | Gene Read Depth | Non-Genic Read Depth | Coverage Ratio |
|---|---|---|---|---|---|---|---|
| PvDBP | LT635617 | 970025-980025 | 980025-988681 | 988681-998681 | 4378.26707 | 1772.59359 | 2.47 |
| PvDBP2 | LT635612 | 94013-104013 | 104013-107429 | 107429-117429 | 2120.53731 | 1800.19293 | 1.18 |
| PvRBP1a | LT635618 | 61106-71106 | 71106-80980 | 80980-90980 | 2397.57195 | 2437.86531 | 0.98 |
| PvRBP1b | LT635618 | 49612-59612 | 59612-70478 | 70478-80478 | 2467.78531 | 2256.64292 | 1.09 |
| PvRBP2a | LT635625 | 102686-112686 | 112686-121381 | 121381-131381 | 2260.37328 | 1917.22883 | 1.18 |
| PvRBP2b | LT635619 | 23312-33312 | 33312-43704 | 43704-53704 | 2305.30347 | 1922.30887 | 1.2 |
| PvRBP2c | LT635616 | 1448611-1458611 | 1458611-1467388 | 1467388-1477388 | 1271.57553 | 1645.0383 | 0.77 |

**Supplementary Table 2. Identification of potential gene duplications in DRC *P. vivax* using read depth.** Read depth for each gene was determined for the DRC *P. vivax* sample using the BAM file with optical duplicates removed. Read depth for the non-genic (upstream and downstream) regions and within the gene itself was estimated by Bedtools, and the coverage ratio is the result of dividing the gene read depth by the non-genic read depth.

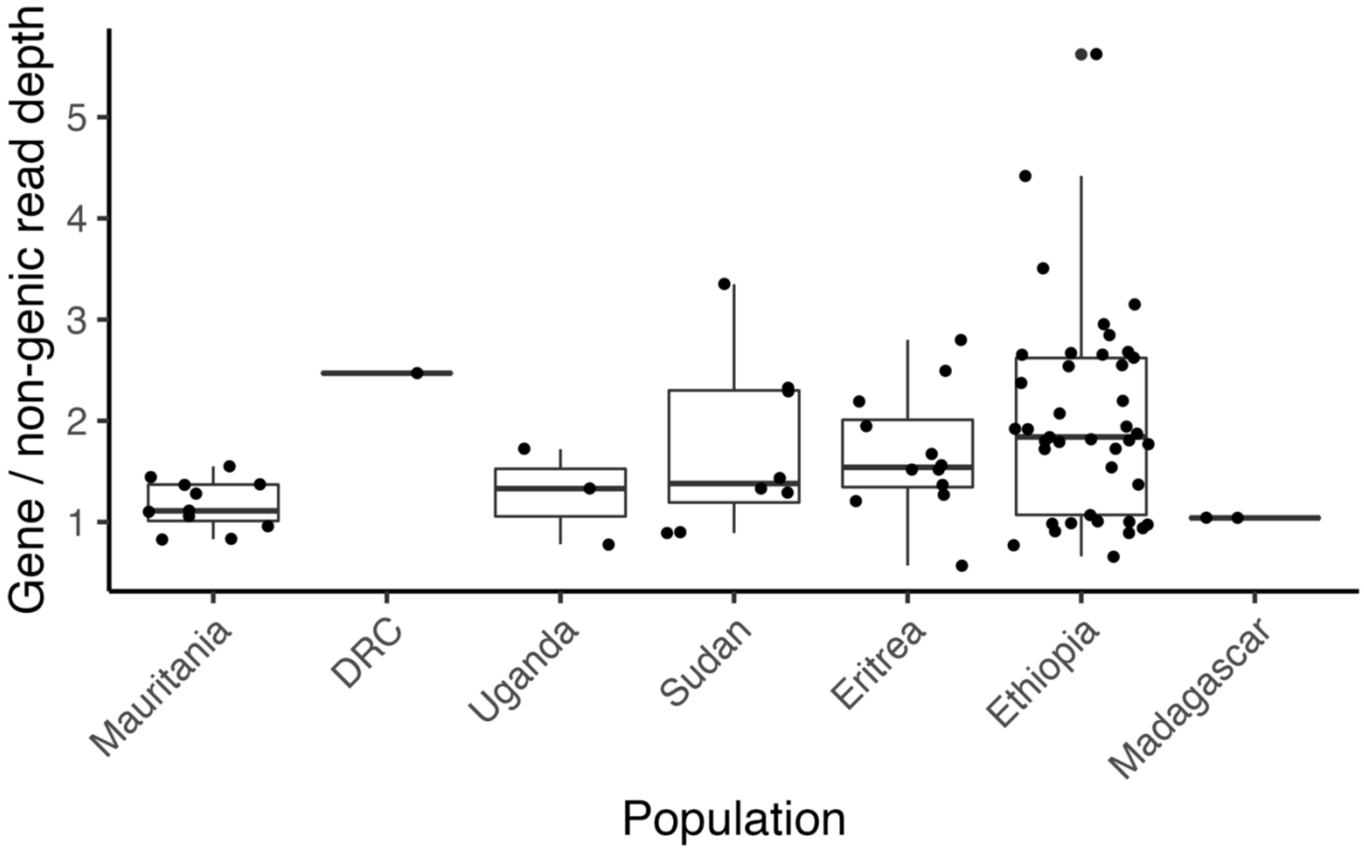


**Supplemental Figure 4. Duplication of PvDBP in African samples**
PvDBP copy number variation for *P. vivax* in each country in Africa. Most countries have a mix of single copy and one or more duplications except for Mauritania and Madagascar.

| Accession | Country | Genic Coverage | Non-Genic Coverage | Coverage Ratio |
|---|---|---|---|---|
| SANRU | DRC | 4378.26707 | 1772.59359 | 2.47 |
| ERR5740704 | Eritrea | 2.14381425 | 1.68838116 | 1.27 |
| ERR5740712 | Eritrea | 7.65415271 | 5.01969803 | 1.52 |
| ERR5740715 | Eritrea | 4.46517269 | 1.59574043 | 2.8 |
| ERR5740716 | Eritrea | 5.67136421 | 3.39906009 | 1.67 |
| ERR5740721 | Eritrea | 0.5459166 | 0.96065394 | 0.57 |
| ERR5740723 | Eritrea | 7.35185399 | 2.94860514 | 2.49 |
| ERR5740726 | Eritrea | 5.63532402 | 3.61338866 | 1.56 |
| ERR5740729 | Eritrea | 5.66270071 | 3.7219778 | 1.52 |
| ERR5740740 | Eritrea | 2.76377498 | 1.26382362 | 2.19 |
| ERR5740742 | Eritrea | 3.27318933 | 1.68043196 | 1.95 |
| ERR5740743 | Eritrea | 2.9459397 | 2.14563544 | 1.37 |
| ERR5740851 | Eritrea | 1.12579416 | 0.92780722 | 1.21 |
| ERR2678989 | Ethiopia | 313.21035 | 70.8569143 | 4.42 |
| ERR2678994 | Ethiopia | 39.3205498 | 40.2654235 | 0.98 |
| ERR2678996 | Ethiopia | 199.716415 | 74.8611639 | 2.67 |
| ERR2678997 | Ethiopia | 14.5378307 | 7.5840416 | 1.92 |
| ERR2678998 | Ethiopia | 65.544877 | 67.9140586 | 0.97 |
| ERR2678999 | Ethiopia | 123.29479 | 68.5157984 | 1.8 |
| ERR2679000 | Ethiopia | 1096.21763 | 194.911109 | 5.62 |
| ERR2679001 | Ethiopia | 30.2270995 | 11.8430157 | 2.55 |
| ERR2679002 | Ethiopia | 306.558739 | 116.855165 | 2.62 |
| ERR2679003 | Ethiopia | 153.339841 | 57.7811719 | 2.65 |
| ERR2679004 | Ethiopia | 1553.42694 | 442.006749 | 3.51 |
| ERR2679005 | Ethiopia | 135.929883 | 135.295171 | 1 |
| ERR2679008 | Ethiopia | 102.424165 | 32.4782522 | 3.15 |
| ERR2679009 | Ethiopia | 98.6847638 | 37.2579242 | 2.65 |
| ERR2679012 | Ethiopia | 33.4857341 | 17.4821018 | 1.92 |
| ERR5740701 | Ethiopia | 2.13029918 | 0.96965304 | 2.2 |
| ERR5740702 | Ethiopia | 2.95749105 | 1.91580842 | 1.54 |
| ERR5740709 | Ethiopia | 1.55307843 | 1.71157884 | 0.91 |
| ERR5740710 | Ethiopia | 1.02968696 | 1.56709329 | 0.66 |
| ERR5740727 | Ethiopia | 9.57560356 | 4.04529547 | 2.37 |
| ERR775189 | Ethiopia | 50.1778907 | 24.2323768 | 2.07 |
| ERR775190 | Ethiopia | 24.7500289 | 13.5826917 | 1.82 |
| ERR775191 | Ethiopia | 123.571214 | 67.3077192 | 1.84 |
| ERR925409 | Ethiopia | 77.1175927 | 27.0638436 | 2.85 |
| ERR925411 | Ethiopia | 53.2547072 | 28.5268473 | 1.87 |
| ERR925412 | Ethiopia | 45.92688 | 25.3045196 | 1.81 |
| ERR925416 | Ethiopia | 18.8858727 | 17.7147285 | 1.07 |

| | | | | |
|---|---|---|---|---|
| ERR925417 | Ethiopia | 42.360633 | 21.8155185 | 1.94 |
| ERR925421 | Ethiopia | 8.97031304 | 9.49540046 | 0.94 |
| ERR925424 | Ethiopia | 19.300104 | 10.9335566 | 1.77 |
| ERR925431 | Ethiopia | 45.0070463 | 26.1352365 | 1.72 |
| ERR925433 | Ethiopia | 88.5899272 | 34.8744626 | 2.54 |
| ERR925435 | Ethiopia | 36.4942821 | 21.1715329 | 1.72 |
| ERR925436 | Ethiopia | 35.6503408 | 19.9080092 | 1.79 |
| ERR925438 | Ethiopia | 103.39136 | 38.509899 | 2.68 |
| ERR925439 | Ethiopia | 72.5393323 | 24.5691931 | 2.95 |
| ERR925440 | Ethiopia | 27.5119556 | 27.8035197 | 0.99 |
| ERR925441 | Ethiopia | 88.8489084 | 88.2980702 | 1.01 |
| SRR14191981 | Ethiopia | 12.8861037 | 9.39046095 | 1.37 |
| SRR14191982 | Ethiopia | 15.5872704 | 20.3619138 | 0.77 |
| SRR14191983 | Ethiopia | 6.57179161 | 7.36591341 | 0.89 |
| ERR490350 | Madagascar | 134.642717 | 130.056644 | 1.04 |
| SRR570031 | Madagascar | 513.619152 | 495.763424 | 1.04 |
| SRR14191966 | Mauritania | 19.6048285 | 12.6476852 | 1.55 |
| SRR14191967 | Mauritania | 14.3873166 | 15.019948 | 0.96 |
| SRR14191970 | Mauritania | 35.0400832 | 24.3187681 | 1.44 |
| SRR14191972 | Mauritania | 10.5723692 | 12.7894211 | 0.83 |
| SRR14191975 | Mauritania | 31.9665011 | 23.3059694 | 1.37 |
| SRR14191976 | Mauritania | 10.4561626 | 9.3930107 | 1.11 |
| SRR14191977 | Mauritania | 20.7820261 | 19.5213479 | 1.06 |
| SRR14191978 | Mauritania | 8.51957953 | 10.3180682 | 0.83 |
| SRR14191979 | Mauritania | 39.9512533 | 31.3178182 | 1.28 |
| SRR14191980 | Mauritania | 34.549151 | 25.2717228 | 1.37 |
| SRR332413 | Mauritania | 520.312579 | 472.131287 | 1.1 |
| ERR5740714 | Sudan | 2.57560356 | 1.92985701 | 1.33 |
| ERR5740717 | Sudan | 4.09587617 | 4.55479452 | 0.9 |
| ERR5740718 | Sudan | 5.42093104 | 2.32651735 | 2.33 |
| ERR5740728 | Sudan | 10.1383851 | 3.02189781 | 3.35 |
| ERR5740852 | Sudan | 1.72115051 | 1.93705629 | 0.89 |
| SRR14191963 | Sudan | 11.8701629 | 9.21417858 | 1.29 |
| SRR14191964 | Sudan | 20.3321012 | 8.8739626 | 2.29 |
| SRR14191965 | Sudan | 17.4111124 | 12.1355864 | 1.43 |
| ERR5740703 | Uganda | 3.10511725 | 2.33686631 | 1.33 |
| ERR5740705 | Uganda | 0.97400947 | 1.25312469 | 0.78 |
| ERR5740706 | Uganda | 2.90666513 | 1.6850315 | 1.72 |

**Supplementary Table 3. PvDBP coverage for all African countries used to generate figure 4B.** See Supplementary table 2 for PvDBP coordinates used.

| pop1 | pop2 | pop3 | pop4 | est | se | z | p |
|---|---|---|---|---|---|---|---|
| pvl | DRC | PNG | Uganda | 0.0031888 | 1.2772E-04 | 24.9661983 | 1.424E-137 |
| pvl | DRC | PNG | Ethiopia | 0.00280492 | 1.4810E-04 | 18.93982009 | 5.358E-80 |
| pvl | DRC | PNG | Madagascar | 0.00274388 | 1.1186E-04 | 24.53023864 | 7.030E-133 |
| pvl | DRC | PNG | Eritrea | 0.00257469 | 1.2403E-04 | 20.75865454 | 1.024E-95 |
| pvl | DRC | PNG | Pakistan | 0.00252949 | 1.2462E-04 | 20.29780614 | 1.345E-91 |
| pvl | DRC | PNG | Afghanistan | 0.00251222 | 1.2886E-04 | 19.49594722 | 1.188E-84 |
| pvl | DRC | PNG | SriLanka | 0.00249463 | 1.4271E-04 | 17.48048597 | 2.018E-68 |
| pvl | DRC | PNG | Bangladesh | 0.00249387 | 1.4986E-04 | 16.6413591 | 3.496E-62 |
| pvl | DRC | PNG | India | 0.00248823 | 1.2973E-04 | 19.18062832 | 5.373E-82 |
| pvl | DRC | PNG | Sudan | 0.00237983 | 1.3586E-04 | 17.51681328 | 1.066E-68 |
| pvl | DRC | PNG | Guyana | 9.85E-04 | 1.0713E-04 | 9.198170239 | 3.641E-20 |
| pvl | DRC | PNG | Brazil | 9.42E-04 | 9.2782E-05 | 10.1498476 | 3.319E-24 |
| pvl | DRC | PNG | Nicaragua | 9.41E-04 | 9.2708E-05 | 10.15543284 | 3.134E-24 |
| pvl | DRC | PNG | Colombia | 9.32E-04 | 1.0243E-04 | 9.101371202 | 8.920E-20 |
| pvl | DRC | PNG | Thailand | 9.27E-04 | 7.5619E-05 | 12.26356511 | 1.421E-34 |
| pvl | DRC | PNG | Mauritania | 9.00E-04 | 1.0978E-04 | 8.196586144 | 2.473E-16 |
| pvl | DRC | PNG | Peru | 8.84E-04 | 1.0074E-04 | 8.773900568 | 1.726E-18 |
| pvl | DRC | PNG | Panama | 8.26E-04 | 1.0662E-04 | 7.74650634 | 9.446E-15 |
| pvl | DRC | PNG | Myanmar | 8.07E-04 | 7.0737E-05 | 11.40500208 | 3.948E-30 |
| pvl | DRC | PNG | Mexico | 8.06E-04 | 1.0881E-04 | 7.404705339 | 1.314E-13 |
| pvl | DRC | PNG | China | 7.40E-04 | 7.3768E-05 | 10.03492804 | 1.070E-23 |
| pvl | DRC | PNG | Cambodia | 5.88E-04 | 6.4353E-05 | 9.133260076 | 6.647E-20 |
| pvl | DRC | PNG | Laos | 5.81E-04 | 9.3595E-05 | 6.212136475 | 5.227E-10 |
| pvl | DRC | PNG | Malaysia | 5.16E-04 | 7.5149E-05 | 6.869492877 | 6.443E-12 |
| pvl | DRC | PNG | Vietnam | 5.07E-04 | 6.4147E-05 | 7.906284601 | 2.652E-15 |
| pvl | DRC | PNG | Indonesia | 2.74E-04 | 3.7911E-05 | 7.223936941 | 5.050E-13 |
| pvl | DRC | PNG | Philippines | 2.22E-04 | 9.6136E-05 | 2.313679263 | 2.069E-02 |
| pvl | DRC | PNG | NorthKorea | 1.36E-04 | 8.5669E-05 | 1.589632817 | 1.119E-01 |

**Supplementary Table 4. F4 statistics calculated using Admixtools2.**
